# A new paradigm for Prelamin A proteolytic processing by ZMPSTE24: the upstream SY^LL cleavage occurs first and there is no CaaX processing by ZMPSTE24

**DOI:** 10.1101/2020.05.13.093849

**Authors:** Laiyin Nie, Eric Spear, Timothy D. Babatz, Andrew Quigley, Yin Yao Dong, Amy Chu, Bérénice Rotty, Rod Chalk, Shubhashish M.M. Mukhopadhyay, Nicola A. Burgess-Brown, Ashley C.W. Pike, Stephen G. Young, Susan Michaelis, Elisabeth P. Carpenter

**Affiliations:** Structural Genomics Consortium, University of Oxford, Old Road Campus Research Building, Roosevelt Drive, Oxford, OX3 7DQ, UK; Department of Cell Biology, The Johns Hopkins University School of Medicine, 725 N Wolfe St., Baltimore, MD 21205, U.S.A; Departments of Medicine and Human Genetics, David Geffen School of Medicine, University of California, Los Angeles, CA, USA

## Abstract

Human ZMPSTE24, an integral membrane zinc metalloprotease, is required for conversion of prelamin A to mature lamin A, a component of the nuclear lamina and failure of this processing causes premature ageing disorders. ZMPSTE24 has also been implicated in both type 2 diabetes mellitus and in viral-host response mechanisms, but to date its only confirmed substrate is the precursor for lamin A. Prelamin A is thought to undergo four C-terminal post-translational modifications in the following order: farnesylation, SIM tripeptide cleavage, carboxymethylation and upstream “SY^LL” cleavage. Here we present evidence that the sequence of events does not follow the accepted dogma. We assessed cleavage of long human prelamin A sequence peptides by purified human ZMPSTE24 combined with FRET and mass spectrometry to detect products. Surprisingly, we found that the “SY^LL” cleavage occurs before and independent of the C-terminal CSIM modifications. We also found that ZMPSTE24 does not perform the predicted C^SIM tripeptide cleavage, but rather it removes an IM dipeptide. ZMPSTE24 can perform a tripeptide cleavage with a canonical CaaX box (C: cysteine; a: aliphatic; X: any residue), but the C-terminus of prelamin A is not a true CaaX sequence. Regardless of the C-terminal modifications of prelamin A, ZMPSTE24 can perform upstream SY^LL cleavage, thus removing the unwanted farnesylated C-terminus. Therefore, it is failure of SY^LL cleavage, not the C-terminal processing that is the likely cause of progeroid disorders.

## Introduction

ZMPSTE24 (also called farnesylated protein-converting enzyme 1, FACE1), is an integral membrane zinc metalloprotease that is required for conversion of prelamin A to mature lamin A, a major protein component of the nuclear intermediate filaments known as lamina [1, 2]. ZMPSTE24 has also been implicated in unclogging the translocon in the endoplasmic reticulum [3], removing human islet amyloid polypeptide (IAPP) from the pancreas in patients with type 2 diabetes mellitus [4], and host defenses against viruses [5, 6].

Despite suggestions that ZMPSTE24 could be involved in diverse functions in cell biology, thus far the only confirmed substrate for ZMPSTE24 is prelamin A [7]. Lamins A, B1, B2, and C form the nuclear lamina, an intermediate filament meshwork that lines the nuclear side of the inner nuclear membrane [8, 9]. The nuclear lamin proteins interact with genomic DNA, the nuclear membrane and nucleoplasmic proteins, and these interactions are thought to be important for an array of biological processes [1, 7–9].

Failure of ZMPSTE24-mediated prelamin A processing, due either to missense mutations in *ZMPSTE24* or prelamin A defects that interfere with ZMPSTE24-mediated processing, result in progeria-like disease phenotypes due to an accumulation of farnesylated prelamin A [10]. Hutchinson-Gilford progeria syndrome (HGPS), mandibuloacral dysplasia type B (MAD-B), and restrictive dermopathy (RD) all involve mutations that interfere with the conversion of farnesylated prelamin A to mature lamin A [11–21]. ZMPSTE24-deficient mice exhibit a striking accumulation of farnesylated prelamin A and exhibit disease phenotypes that resemble those in the aforementioned progeroid disorders, including severe growth retardation, nonhealing bone fractures, alopecia, and progressive inanition [1, 2]. Studies of fibroblasts from affected patients have suggested that the severity of disease phenotypes in patients is proportional to the accumulation of farnesylated prelamin A proteins in cells [22–25]. There is some evidence to suggest that defective prelamin A processing could be relevant to physiological aging, at least in certain tissues [26, 27]. Given the relevance of prelamin A processing for health and disease, we believe that it is important to have a clear understanding of the biochemistry of prelamin A processing.

ZMPSTE24 is required for posttranslational processing at the C-terminus of prelamin A, a 661-residue protein that includes a C-terminal CSIM sequence. The textbook view of prelamin A processing is that prelamin A undergoes a series of four sequential posttranslational modifications, including (i) farnesylation of the cysteine by protein farnesyltransferase, (ii) removal of the C-terminal –SIM tripeptide by either RCE1 or ZMPSTE24, (iii) carboxymethylation of the newly exposed farnesylcysteine by ICMT, and (iv) a final endoproteolytic processing step at an SY^LL site located 15 residues upstream from the C-terminal farnesylcysteine methyl ester (S1 Fig). The first three proposed steps are identical to those that occur in a large number of proteins with a C-terminal CaaX motif (where C is cysteine, a is an aliphatic residue and X is any residue) [28], including the Ras GTPases [29]. All such proteins undergo protein farnesylation, release of the C-terminal tripeptide by RCE1, and carboxymethylation of the farnesylcysteine by ICMT, enzymatic reactions collectively known as the CaaX processing reactions. However, no CaaX protein, apart from prelamin A, undergoes an additional C-terminal cleavage reaction, so they all retain their C-terminal farnesylcysteine methyl ester.

Unlike most proteins harboring a classic CaaX motif, prelamin A undergoes a second endoproteolytic cleavage step, catalyzed by ZMPSTE24, at a SY^LL motif located 18 residues from the C-terminus. This cleavage step removes the farnesylcysteine (fCys) from the protein and releases mature lamin A (S1 Fig) [7, 30]. It is generally accepted that prelamin A must undergo all of the CaaX processing steps before the SY^LL cleavage can occur, although two reports raised the possibility that the SY^LL cleavage could occur in the absence of CaaX processing [31, 32].

Many of the studies on the function of the human ZMPSTE24 have been based on comparisons with the yeast orthologue of ZMPSTE24, Ste24, the only other protein thought to perform both C^aaX and upstream proteolysis steps [33]. ZMPSTE24 and Ste24 share an unusual protein fold [34, 35], with seven transmembrane α-helices forming an α-helical barrel enclosing a large, water-filled chamber inside the membrane. The zinc metalloprotease domain lies on the nucleoplasmic or cytoplasmic side of membrane, with the active site facing into the chamber.

Not only are the structures of ZMPSTE24 and Ste24 similar, they also appeared to have very similar proteolytic activities. In yeast Ste24 is required for the biogenesis of the mating pheromone **a**-factor (38). **A**-factor undergoes the classic CaaX processing steps (farnesylation of the C-terminal cysteine, C^aaX tripeptide cleavage, carboxymethylation of the farnesylcysteine) followed by two upstream cleavage reactions. Ste24 and Rce1 can both catalyze the removal of the tripeptide from yeast **a** factor’s classic CaaX motif (CVIA). Then Ste24 catalyses the first of two upstream cleavage reactions, the second being catalysed by Axl1 [33]. Human ZMPSTE24 complements the defect in **a**-factor production in yeast lacking both Rce1 and Ste24, indicating that ZMPSTE24 is capable of cleaving both the C^aaX site in **a**-factor and the upstream cleavage reaction [7, 33], Although some evidence suggests CaaX cleavage may not be a prerequisite for second Ste24-mediatd cleavage of the a-factor precursor [36], it is generally thought that the a-factor C^aaX cleavage occurs before the upstream cleavage and that the C^aaX cleavage is necessary for the upstream cleavage to occur.

Given these observations in the yeast system, it seemed reasonable to assume that ZMPSTE24 would perform both endoproteolytic processing steps on its natural substrate, prelamin A. Thus far, however, there has been no direct experimental evidence to confirm that ZMPSTE24 does cleave the CSIM sequence of prelamin A at the predicted C^aaX site. In fact, when we examined ZMPSTE24-mediated processing of a short farnesylated prelamin A peptide (QSPQNC(f)SIM) with a mass spectrometry– based assay, we found release of the dipeptide IM, raising the possibility of a CS^IM cleavage rather than a C^SIM cleavage. Moreover, the complex that we observed between ZMPSTE24 and a CSIM peptide in the crystal structure suggested that the peptide might undergo a CS^IM cleavage. More studies were needed to explore these preliminary observations based on short peptides.

The biochemistry of the SY^LL cleavage in prelamin A is also relatively unexplored. There is little data available on the kinetics of proteolysis of prelamin A based substrates by ZMPSTE24 or comparison of the SY^LL cleavage, which is unique to ZMPSTE24, and the CSIM cleavage, which can also be performed by RCE1. Available studies followed the appearance of the final lamin A product and/or whether it was possible to carboxymethylate the C-terminus [28, 35, 37–43]. Studies involving purified ZMPSTE24 characterized the C^aaX or upstream reaction using a short peptide based on the C-terminal sequence of human K-RAS protein [44] or the yeast **a**-factor peptide [43] rather than the actual prelamin A sequence. Another study with purified ZMPSTE24 used in native mass spectrometry to investigate complexes of ZMPSTE24 with peptide substrates and products with the SY^LL site but not the CSIM site [45].

In this study, our goal was to provide a biochemical characterization of prelamin A processing by ZMPSTE24, including kinetics for each cleavage reaction (CSIM and SYLL) and identification of the products formed by each processing step. We used peptides containing only one of the two sites, as well as a longer prelamin A peptides containing both cleavage sites. Our results confirmed the dipeptide cleavage from the CSIM site. More importantly, we investigated the relationship between the SYLL and CSIM cleavage reactions and discovered that the processing of prelamin A by ZMPSTE24 does not conform to textbook dogma, which holds that the upstream cleavage reaction at the SYLL site can only occur after the processing at the CSIM motif is complete. Here, we show that ZMPSTE24 performs the upstream SY^LL cleavage reaction in prelamin A, regardless of any post-prenylation processing at the CSIM motif.

## Results

### ZMPSTE24 reaction kinetics suggests faster cleavage at the SY^LL than CSIM site

ZMPSTE24 cleaves prelamin A at two sites (SYLL and CSIM). To measure the kinetics of ZMPSTE24-mediated cleavage reactions, we designed a fluorescence resonance energy transfer (FRET) assay using two farnesylated prelamin A peptides, a 20-mer peptide covering the SYLL site (Peptide 1, Table 1, Fig 1A) and a 9-mer peptide covering the C-terminal CSIM site (Peptide 2, Table 1, Fig 1A). The peptides were modified with a dinitrophenyl (Dnp) and n-amino benzoic acid (Abz) groups on either side of the cleavage site. In the absence of peptide cleavage, the Dnp group quenches the fluorescent signal from the Abz group (Fig 1B). Cleavage of the peptide separates the fluorophore and quencher, resulting in an increase in the fluorescence signal.

**Table 1.** Sequences of C-terminus of human prelamin A and the synthetic peptides used in this study.

| Name | Sequence |
| --- | --- |
| Prelamin A | SFGDNLVTR**SYLL**GNSSPRTQSPQNC( **f** )**SIM** |
| Peptide 1 | ( **Abz** )VTR**SYLL**GK( **Dnp** )SSPRTQSPQNC( **f** ) |
| Peptide 2 | ( **Abz** )QSPSNC( **f** )**SIK**( **Dnp** ) |
| Peptide 3 | LLGNSSPRTK( **Dnp** )SPQNC( **f** )**SIK**( **Abz** ) |
| Peptide 4 | SFGDNLVTR**SYLL**GNSSPRTQSPQNC( **f** )**SIM** |
| Peptide 5 | VTR**SYLL**NGSSPRTQSPQNC( **f** )**VLS** |
| Peptide 6 | SFGDNLVTR**SYLL**GNSSPRTQSPQNC**SIM** |
| Abz, 2-aminobenzoyl group (green); K(Dnp), lysine with a dinitrophenyl modification (red).<br>Peptide sequences are aligned with the sequence of human prelamins A; cleavage sites are depicted with bold font. |  |

**Fig. 1.**
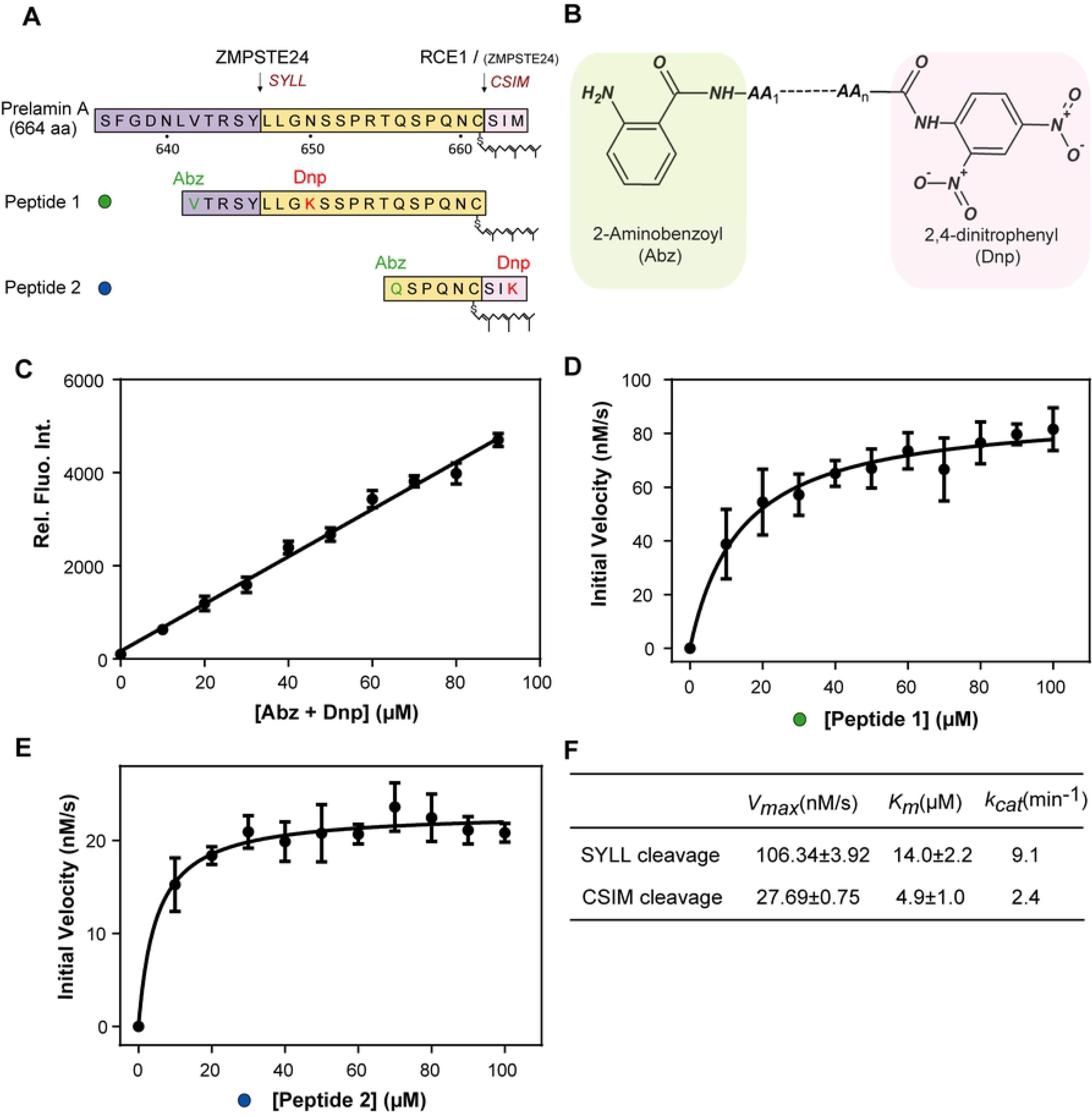
Kinetics of the prelamin A peptide cleavage by ZMPSTE24, as judged by a fluorescence resonance energy transfer (FRET) assay. (A) Schematic representation of fluorescent prelamin A peptides, corresponding to sequences within the carboxyl-terminal region of prelamin A. Green and red indicate attachment sites for a 2-aminobenzoyl (Abz) group and a 2, 4-dinitrophenyl (Dnp) group, respectively. (B) Chemical structures of Abz and Dnp. (C) Correlation between relative fluorescence intensity and peptide concentration. (D–E) Michaelis-Menten saturation curves for ZMPSTE24-mediated cleavage of (D) Peptide 1, representing the SYLL reaction and (E) Peptide 2, representing the CSIM reaction. Data show the mean ± S.D. of three biological replicates, each with three technical repeats. The curve was fitted with this linear regression equation: y = 50.75x + 167.4; R^2^ = 0.992. (F) Kinetic parameters for the two cleavage reactions. The S.E. values represent the error with respect to curve fitting.

Wild-type (WT) human ZMPSTE24 and an active-site ZMPSTE24 mutant (E336A) were purified in n-dodecyl β-D-maltoside (DDM) and incubated with increasing concentrations of prelamin A peptide substrates [34]. The time course of the cleavage reactions, as judged by the FRET signal, were then recorded (S2A and S2B Fig). For both peptides, there was a time-dependent increase in the intensity of the fluorescent signal, reflecting cleavage of the prelamin A peptide. Incubation of the prelamin A peptides with ZMPSTE24^E336A^, which lacks a crucial catalytic residue, did not yield an increase in the fluorescent signal, establishing the specificity of the FRET assay (S2C and S2D Fig). We produced human ZMPSTE24 in both *Sf9* insect cells and in human Expi293F™ cells and compared the activity of protein produced in these two systems. We found that the activities of ZMPSTE24 obtained from different expression systems was very similar (S2E and S2F Fig), confirming that the results we obtained were not an artefact of using *Sf*9 cells for expression. The conversion factor for the relative fluorescence intensity (RFI) and moles of the peptide cleavage product was determined with a standard curve (Fig 1C). The fitted line plot (R^2^ = 0.992) revealed that the linear regression followed the experimental data almost exactly, indicating that intermolecular quenching of the fluorescent signal was negligible.

Next, we compared kinetic parameters for the two cleavage reactions. Initial velocities for peptide 1 cleavage at the SYLL site and the peptide 2 cleavage at the CSIM site were plotted against substrate concentrations and fitted to the Michaelis-Menten equation (Fig 1D and 1E). The kinetic parameters (*K_m_*, *V_max_* and *k_cat_*) are shown in Fig 1F. The substrate affinity was in the low micromolar range for both reactions but was slightly higher affinity with the CSIM reaction. On the other hand, the turnover of the SYLL peptide was nearly five-fold higher than the CSIM peptide, indicating that the SYLL cleavage is faster.

### SYLL peptide cleavage occurs at the expected location, but the CSIM peptide cleavage gives a dipeptide product

We used electrospray ionization mass spectrometry to characterize the peptide cleavage products of the reactions described above (Fig 2 and 3). For peptide 1, which spans the SYLL site, two products were detected: an N-terminal 5-mer peptide (Peptide 1-1) and a C-terminal 15-mer peptide (Peptide 1-2), which eluted at 1.8 and 6.3 min, respectively (Fig 2C). The identity of each peptide was confirmed by mass spectrometry (Fig 2D and 2E). As predicted, the cleavage occurred between Y and L. After a 30-min incubation with ZMPSTE24 at 37°C, the substrate peptide (Peptide 1) was almost completely consumed, with only a small peak remaining at 1356.65 remaining (6.6 min peak in Fig 2C and 2F). We also observed a dimer of the detergent DDM, which co-eluted with peptides 1 and 1-2. This DDM species has been observed previously in mass spectrometry–based analyses of membrane proteins [46].

**Fig. 2.**
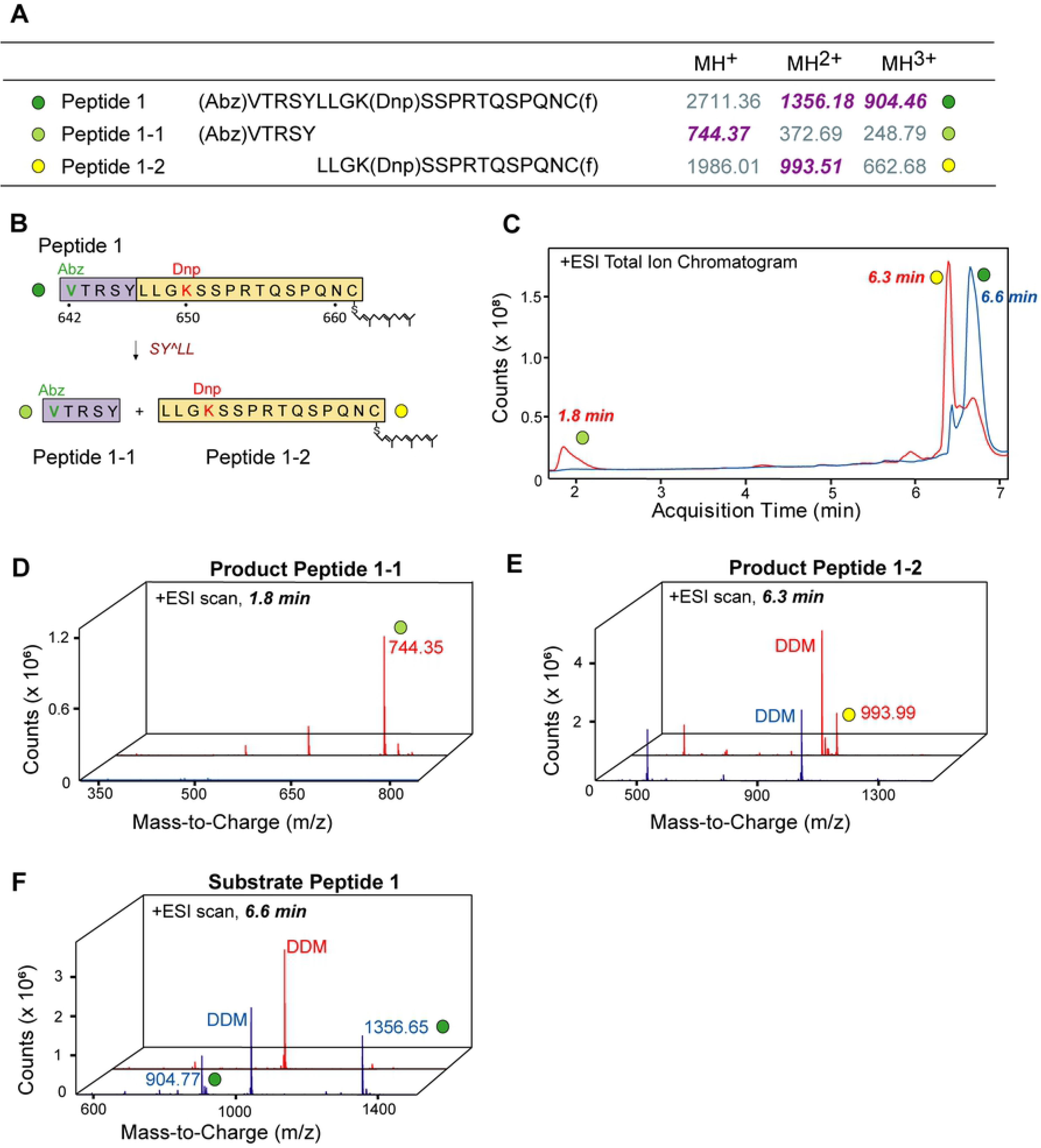
Mass spectrometric analyses of the peptide products from the SYLL cleavage reaction. Peptide 1 was incubated for 30 min at 37°C, with or without ZMPSTE24; the reaction products were separated by HPLC and analyzed by mass spectrometry. (A) Theoretical m/z values of the substrate peptide and the reaction products. M/z values detected are depicted in purple bold italicized font; products that were not detected are shown in grey. (B) Schematic representation of peptide 1 and its cleavage products (colours are the same as in Fig. 1). (C) Total ion chromatograms (TIC) for peptide 1, after incubating the peptide for 30 min at 37°C in the presence (red) and absence (blue) of ZMPSTE24. (D-F) Mass spectrometric spectrum for peptide 1 and reaction products at the indicated time points in the presence (red) or absence (blue) of ZMPSTE24. The retention time for peptide 1 was 6.6 min; the reaction products eluted at 1.8 and 6.3 min C-F. The peptide substrate and the reaction products are marked with coloured dots. The detergent DDM (a DDM dimer with an Na^+^ adduct) was detected in 6.3- and 6.6-min scans. Four biological replicates were obtained; all yielded similar results.

**Fig 3.**
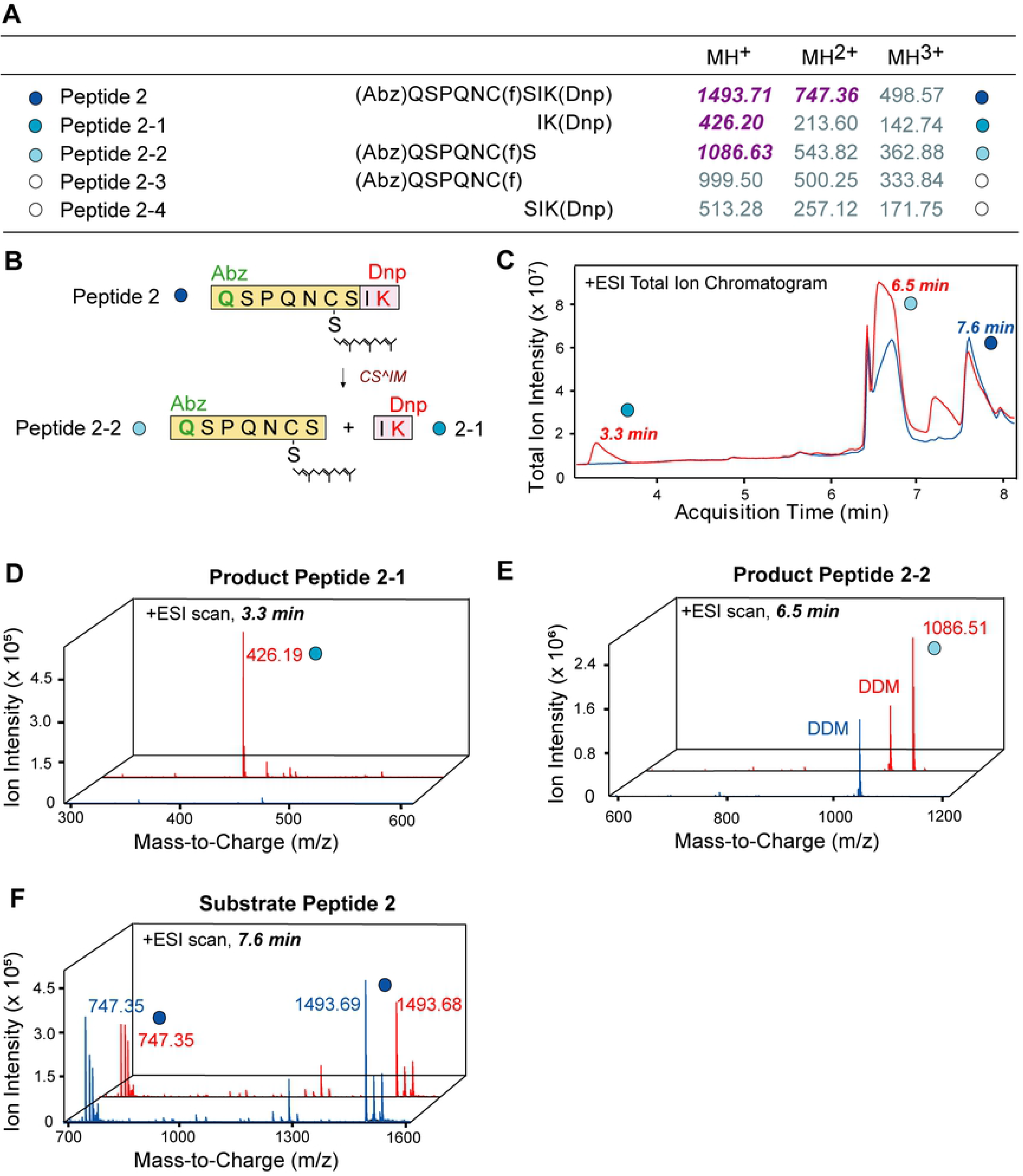
Mass spectrometric analyses of the peptide products of the CSIM cleavage reaction. Peptide 2 was incubated for 30 min at 37°C, with or without ZMPSTE24, the reaction products were separated by HPLC and analysed by mass spectrometry. (A) Theoretical m/z values for the peptide substrate and the reaction products (coloured as in Fig 2). (B) Schematic representation of peptide 2 and its cleavage products (coloured as in Fig 1). (C) Total ion chromatogram (TIC) for peptide 2 in the presence (red) and absence (blue) of ZMPSTE24. (D-F) Mass spectrometric spectrum for peptide 1 and the reaction products at the indicated time points in the presence (red) or absence (blue) of ZMPSTE24. The retention time of the peptide substrate was 7.6 min; the retention times for the two reaction products were 3.3 and 6.5 min (C-F). The substrate and reaction products are marked with coloured dots; peptides that were not detected are marked with white dots. DDM was detected in the product sample at 6.5 min. This experiment was repeated four times, with similar results each time.

To investigate the CSIM cleavage, we analyzed the products from treatment of the farnsylated C-terminal 8-mer peptide (peptide 2) with ZMPSTE24 (Fig 3). We detected two products: peptides 2-1 and 2-2 (Fig 3C-F). Interestingly, neither of those peptides corresponded to the reactions products expected from a classic C^aaX cleavage. Human ZMPSTE24 is thought to function as a prelamin A CaaX protease because the yeast orthologue, Ste24, serves as a CaaX protease, removing a tripeptide from the CaaX motif (CVIA) of yeast **a**-factor [33, 47]. Therefore, we expected that human ZMPSTE24 would cleave peptide 2 between the farnesylcysteine and adjacent serine within the C^SIM motif (a reaction that would release peptides 2-3 and 2-4). However, the products we identified by mass spectrometry were peptides 2-1 and 2-2 (Fig 3C-F), indicating that the cleavage actually occurs between the serine and the isoleucine of the CSIM motif (CS^IM). These results are consistent with our previous observation that with a nonderivatized farnesylated peptide (QSPQNC(f)SIM), ZMPSTE24 removed an IM dipeptide [34]. This finding is also consistent with our structure of a CSIM tetrapeptide in complex with ZMPSTE24, where the tetrapeptide appeared to be positioned (with respect to the Zn^2+^ ion) for a CS^IM cleavage reaction [34]. In the case of peptide 2, most of the substrate was uncleaved after a 30-min incubation with ZMPSTE24 (Fig 3F), indicating that the CS^IM cleavage reaction was far slower than the SY^LL reaction.

To determine if there were any reaction products consistent with a C^SIM cleavage, we extracted ion chromatograms at specific m/z values for peptides 2-3 and 2-4. Small peaks were observed, and the ESI spectrum confirmed the existence of both peptides (S3B, S3C, S3E and S3F Fig). However, the signal intensities for the extracted ion chromatogram and the corresponding ESI spectrum for peptides 2-3 and 2-4 were 100-folder lower than for the dipeptide cleavage products (Peptides 2-1 and 2-2) (S3A and S3D Fig). Moreover, we only identified peptides 2-3 and 2-4 in two of four biological replicates; in the other two, peaks for peptides 2-3 and 2-4 were undetectable. Therefore, our results suggest that although ZMPSTE24-mediated cleavage of Peptide 2 releases a tripeptide but only in very small amounts; the primary reaction is a CS^IM cleavage, which yields an IM dipeptide product.

It has been reported that after cleavage of a farnesylated CSIM containing peptide by ZMPSTE24, one of the products could be carboxymethylated by ICMT [37]. Since carboxymethylation requires an exposed farnesylcysteine, the product of a tripeptide C^aaX cleavage, this finding suggested that some C^aaX peptide processing had occurred. However, the level of carboxymethylation in those studies was low [37], consistent with the very low levels of the C^aaX cleavage in our mass spectrometry studies. This method would not detect the products of a CS^IM cleavage, since an exposed C(f)S would not be a substrate for ICMT, so it does not reveal how much dipeptide release occurred.

Our studies suggest that a CS^IM dipeptide cleavage is the preferred ZMPSTE24 reaction [34], but those experiments were performed with a short (9-mer) peptide. To exclude the possibility that the use of a short peptide lead to abnormal cleavage, we repeated the FRET based assay with a derivatised 18-mer peptide (Peptide 3) (S4A Fig). The time course activity profile (S4B Fig) indicated that ZMPSTE24 does cleave the longer peptide, although the activity level was low, with a large quantity of the substrate still present after 30 mins (S4E Fig), so it was not possible to accurately measure reaction kinetics. Analysis of the products of the cleavage of the 18-mer peptide by ZMPSTE24 showed mainly formation of a 16-mer product (Peptide 3-1), indicating a dipeptide CS^IM cleavage (S4D Fig). Although there was a small peak for the 15-mer product (Peptide 3-2), it was only 1% of the size of the Peptide 3-1 peak, suggesting that ZMPSTE24 very much favours the CS^IM cleavage (S3A and S3B Fig, S5A and S5B Fig). The observation that the CS^IM dipeptide cleavage occurs with both the 9-mer and 18-mer peptides indicates that the dipeptide cleavage is not the result of use of a shorter peptide.

### ZMPSTE24 performs the SY^LL cleavage before the CS^IM cleavage

Our studies with FRET-labelled peptides revealed that the SY^LL cleavage was fivefold faster than the CSIM cleavage and the CSIM cleavage involved the release of a dipeptide rather than a tripeptide. However, these studies involved peptide substrates modified by FRET labels. To exclude the possibility that the labels affected the cleavage process, we investigated ZMPSTE24 activity against a native 29-mer farnesylated prelamin A peptide containing both cleavage sites (Peptide 4). We used mass spectrometry to assess peptide product formation at multiple time points (Fig 4). If the CS^IM cleavage were to occur before the SY^LL cleavage (as generally assumed), we would expect Peptide 4-5 (or Peptide 4-6) to appear in the reaction mixture first, followed later by the appearance of Peptides 4-1 and 4-4. However, that was not the case.

**Fig 4.**
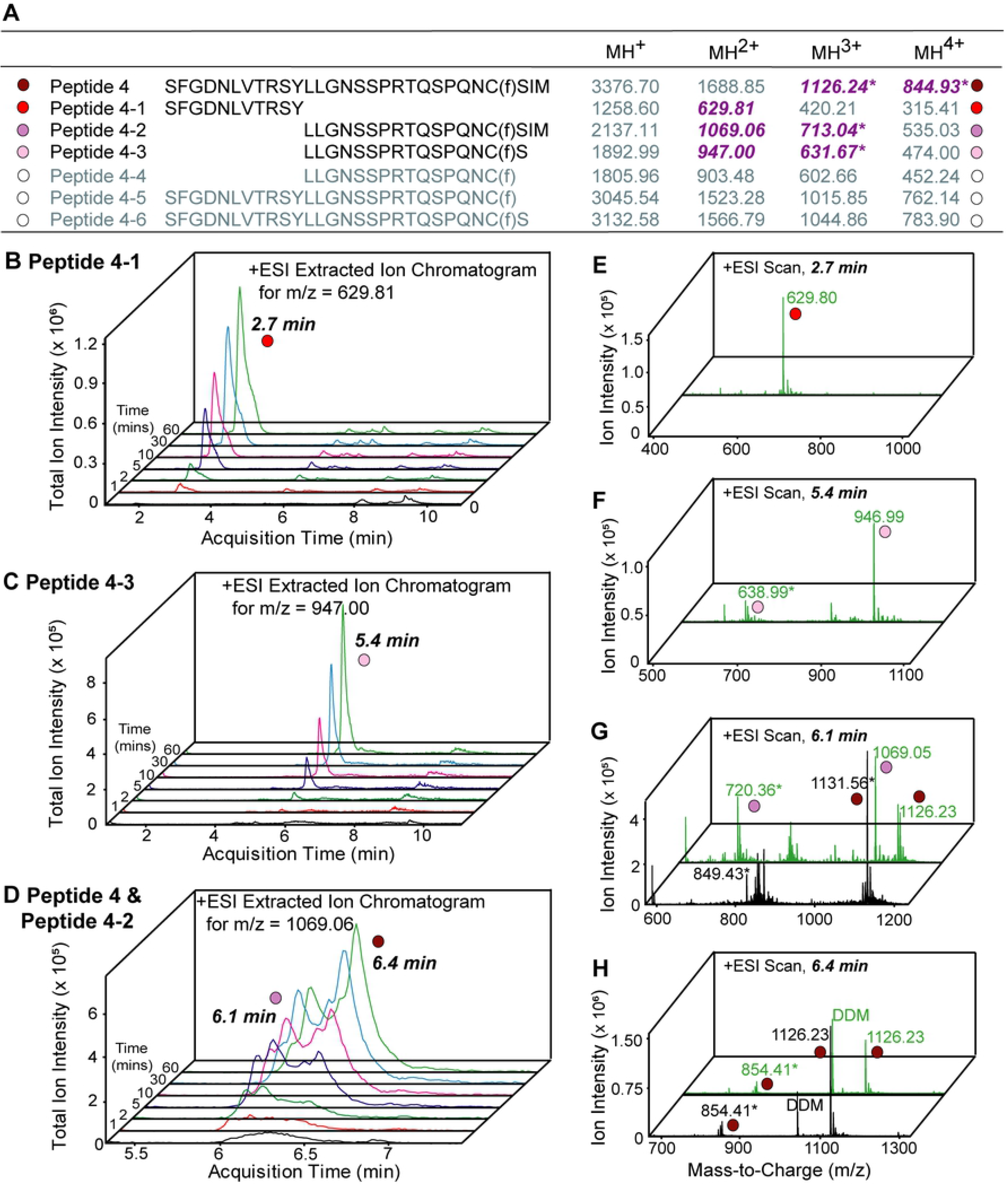
ZMPSTE24-mediated cleavage of a farnesylated 29-mer prelamin A peptide. After incubating Peptide 4 with ZMPSTE24 at 37°C for the specified times, both peptide substrates and products were separated by HPLC and analysed by mass spectrometry. (A) Theoretical m/z values for the substrate and product peptides (colour scheme as in Fig. 1). Peptides with Na^+^ adducts are labelled with an asterisk. (B-D) Extracted ion chromatogram (EIC) for indicated m/z values at time point 0 (black), 1 min (red), 2 min (olive), 5 min (purple), 10 min (magenta), 30 min (blue), and 60 min (green). (E-H) Mass spectra of the 0-min (black) and 60-min (green) samples. The substrate and product peptides detected are marked with coloured dots; peptides that were not detected are marked with white dots. The peptide substrate (peptide 4, red dot) has a retention time of 6.4 min and was detected in triply- and quadropoly-charged form with a Na^+^ adduct (D,H); the product peptide 4-1 (red dot) has a retention time of 2.7 min and was detected in its doubly-charged form (B,E); product peptides 4-2 (lavender dots) and 4-3 (pink dots) eluted at 6.1 and 5.4 min, respectively, and were detected in doubly- and triply-charged forms with Na^+^ adducts (C,D,F,G). DDM co-eluted with peptide 4 at 6.4 min (G). The time-course mass spectrometry measurements were performed in three biological replicates.

Contrary to expectations, the SY^LL cleavage occurred first, releasing Peptides 4-1 and 4-2 (Fig 4A, 4B and 4D). Peptide 4-1 was formed continuously and accumulated progressively (Fig 4B), whereas the amount of peptide 4-2 (the C-terminal product of the SY^LL cleavage) increased briefly but then remained constant (Fig 4D), implying the existence of an equilibrium between production and consumption of the peptide. If there were an initial SY^LL cleavage followed by a CS^IM dipeptide cleavage, peptide 4-3 would be formed. Indeed, peptide 4-3 did appear (Fig 4C) and the amount of that peptide increased from 30 to 60 min. Furthermore, mass spectra of the 1-min sample revealed a greater abundance of the product of an initial SY^LL cleavage reaction (peptide 4-2) than the product resulting from both cleavage reactions (peptide 4-3) (S6A and S6B Fig). We did not detect peptides 4-4 or 4-5 (products that would have formed from a C^aaX tripeptide cleavage), nor did we detect peptide 4-6 (which would have indicated that the CS^IM cleavage occurred before the SY^LL cleavage). Thus, our data indicates that the first reaction is the SY^LL cleavage (yielding Peptide 4-2), which subsequently undergoes a CS^IM cleavage (yielding Peptide 4-3).

We considered the formal possibility that the initial SY^LL cleavage was a peculiarity of ZMPSTE24 expression in *Sf*9 cells. To explore this possibility, we repeated the time course mass spectrometry measurements with ZMPSTE24 that had been purified from HEK293 cells. We obtained similar results, regardless of the expression system used to produce the protein. To exclude artefacts resulting from purification of ZMPSTE24 in detergent micelles, we performed studies with ZMPSTE24 that had been reconstituted into proteoliposomes. Once again, similar results were obtained with protein in micelles or proteoliposomes (S7A-S7D Fig). We observed production of Peptide 4-1 and 4-3 (S7A and S7B Fig) and an equilibrium between the formation and consumption of Peptide 4-3 after 10 min (S7C Fig). Peptide 4-2 still not resolved well because its properties resembled the peptide substrate (S7C and S7D Fig). Thus, the time course experiments revealed that the SY^LL cleavage occurs first, followed by a subsequent CS^IM site (releasing an IM dipeptide).

### A prelamin A peptide with a C-terminal CVLS is cleaved at the SY^LL site first, then undergoes C^VLS cleavage

There are two potential explanations for the CS^IM cleavage by ZMPSTE24—either ZMPSTE24 is not a *bona fide* C^aaX protease and is only capable of the di-peptide cleavage or the CSIM site in prelamin A is not a *bona fide* CaaX motif and cannot undergo a C^aaX cleavage. To address this issue, we performed time course mass spectrometry studies with a 23-mer farnesylated prelamin A peptide (peptide 5) in which the C-terminal CSIM sequence had been replaced with CVLS (the CaaX motif in H-RAS) (Fig 5A). The ion chromatogram revealed a progressive increase in the C-terminal product of the SY^LL cleavage (Peptide 5-1) (Fig 5C). Both doubly- and triply-charged states were detected (with a sodium adduct in the latter) (Fig 5D). If ZMPSTE24 were to cleave within the CVLS site, we would expect to find either peptides 5-2 and/or 5-3, depending on whether a tripeptide or a dipeptide was released. Although the m/z value for the doubly-charged Peptide 5-2 were close to that of the triply-charged Peptide 5, it was possible to resolve the peptides on the extracted ion chromatogram (Fig 5E-5G). Peptide 5-2 was identified, indicating that the tripeptide C^aaX cleavage was favoured (Fig 5F). The peak for peptide 5-2 appeared only after 5 min, whereas the peak for Peptide 5-1 could be detected at 1 min (Fig 5E), indicating that the SY^LL cleavage preceded the C^VLS cleavage. Moreover, the peak for peptide 5-2 was small, reflecting a lower efficiency for the C^VLS reaction than the SY^LL reaction. Thus, a prelamin A peptide terminating with a CVLS motif first undergoes the SY^LL cleavage and subsequently a C^aaX cleavage. These findings suggest that ZMPSTE24 is capable of acting as a *bona fide* C^aaX protease but not with a substrate terminating in CSIM.

**Fig 5.**
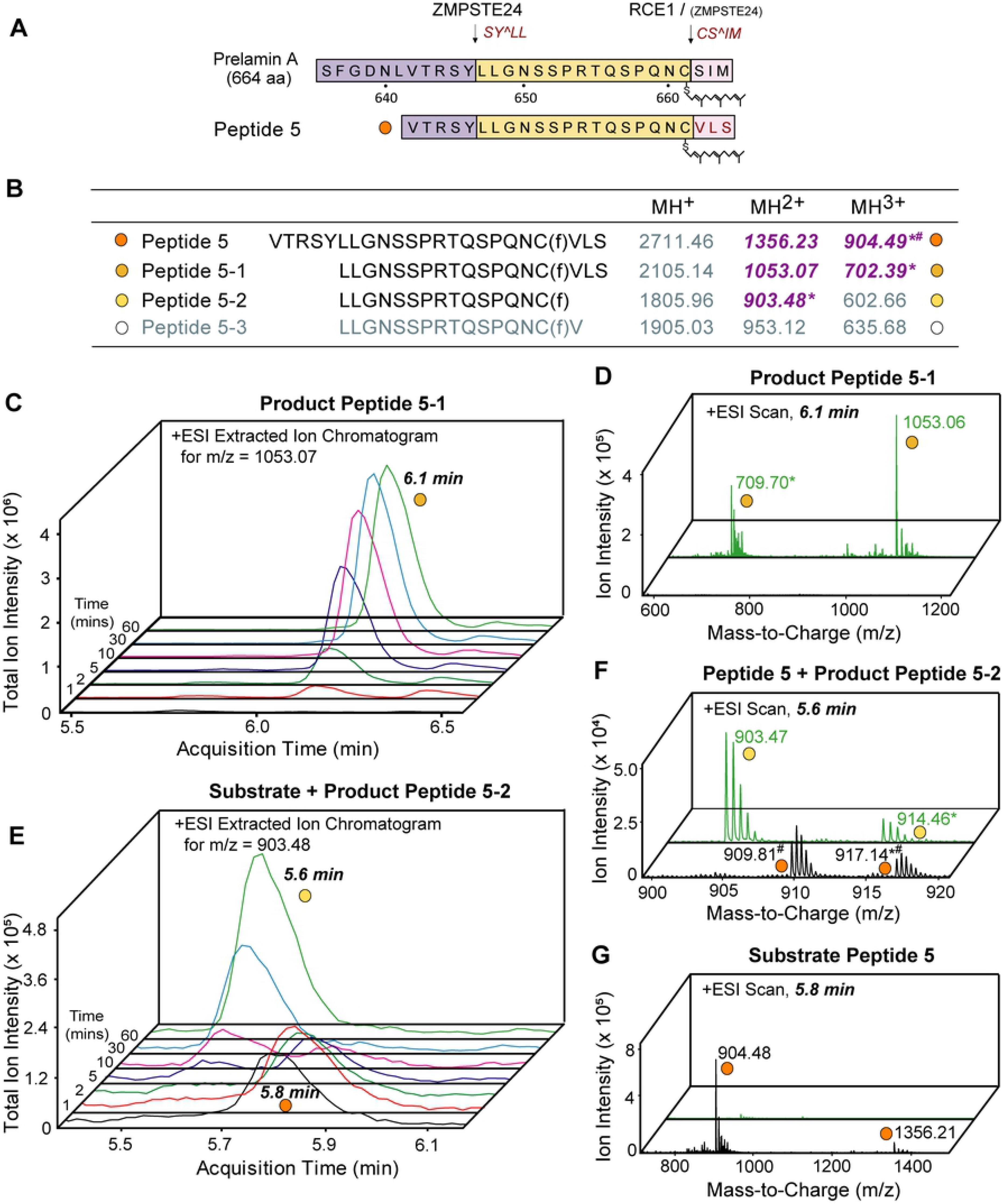
ZMPSTE24-mediated processing of a prelamin A peptide terminating in CVLS involves an initial SY^LL cleavage followed by a C^aaX cleavage. (A) Schematic representation of peptide 5. The modified C-terminal residues are marked in red. (B) Theoretical m/z values for the substrate and product peptides. The peptides that were detected are shown in purple and bold italic font, and those that were not detected are in are in grey; peptides detected with Na^+^ adduct are labelled with an asterisk, and peptides detected in an oxidised form are labelled with a hash. (C-D) Extracted ion chromatogram (EIC) for the m/z values at time point 0 (black), 1 min (red), 2 min (olive), 5 min (purple), 10 min (magenta), 30 min (blue), and 60 min (green). (D,F-G): Mass spectra at 0 (black) and 60 min (green). Substrate and product peptides are marked with coloured dots; peptides that were not detected are marked with white dots. Product peptide 5-1 (peach dot) has a retention time of 6.1 min and was detected in doubly- and triply-charged forms with a Na^+^ adduct (C-D); product peptide 5-2 (yellow dot) has a retention time of 5.6 min and was detected in its doubly-charged form with a Na^+^ adduct (E and F); the peak at 5.8 min was substrate peptide 5 (orange dot) in its doubly- and triply-charged forms (E,G). The mass spectrometry measurements were performed in three biological replicates.

### The farnesyl lipid modification influences the order in which the cleavage reactions occur

To investigate the importance of the farnesyl lipid on ZMPSTE24-mediated prelamin A cleavage, we performed time-course mass spectrometry studies with a nonfarnesylated 29-mer prelamin A peptide (Fig 6). The first processing event was a SY^LL cleavage (Peptides 6-1 and 6-2), but we also observed peptide products resulting from a CS^IM cleavage (Peptides 6-3 and 6-6). We found no evidence for a C^SIM cleavage (Peptides 6-4 and 6-5). The peak corresponding to Peptide 6-6 appeared at approximately the same rate as the SY^LL cleavage products, suggesting that CS^IM cleavage and SY^LL reactions occur simultaneously in the absence of the farnesyl lipid modification, unlike the faster reaction we observed at SY^LL when farnesyl is present (Fig 3).

**Fig 6.**
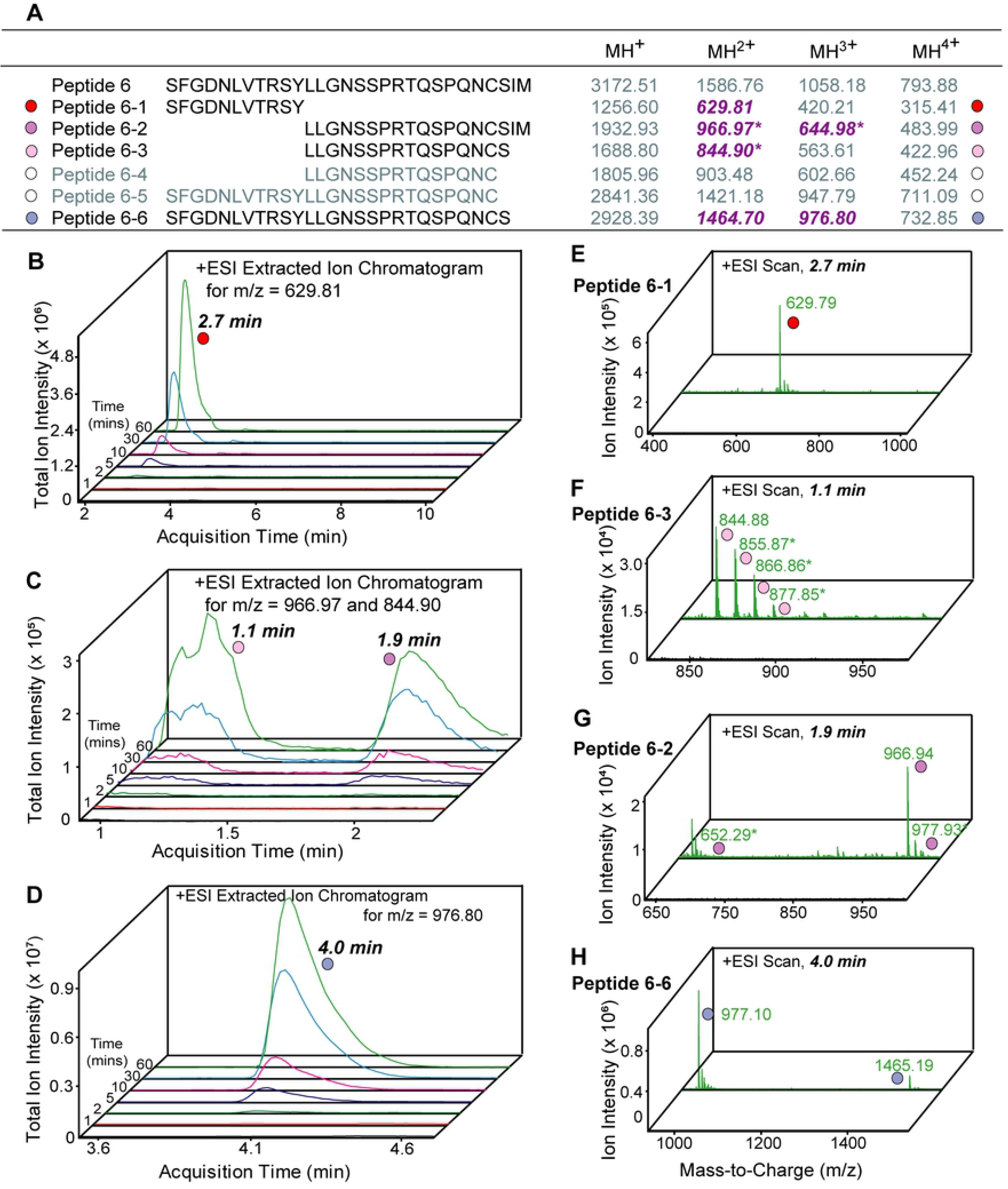
ZMPSTE24-mediated processing of a nonfarnesylated 29-mer prelamin A peptide. (A) Theoretical m/z values for the substrate and product peptides (coloured as in Fig. 1). Peptides detected with an Na^+^ adduct are labelled with an asterisk. (B-D) Extracted ion chromatogram (EIC) for indicated m/z values at 0 (black), 1 min (red), 2 min (olive), 5 min (purple), 10 min (magenta), 30 min (blue), and 60 min (green). (E-H) Mass spectra of 0-(black) and 60-min (green) samples. The substrate and product peptides that were detected are marked with coloured dots, whereas peptides that were not detected are marked with white dots. The Peptide 6-1 product (red dot) has a retention time of 2.7 min and was detected in a doubly-charged form (B,E). Peptides 6-2 (lavender dots) and 6-3 (pink dots) eluted at 1.9 and 1.1 min, respectively and were detected in doubly- and triply-charged forms with a Na^+^ adduct (C,F,G). Peptide 6-6 (periwinkle dot) has a retention time of 4.0 min and was detected in doubly- and triply charged forms. The time course mass spectrometry experiments were performed with three biological replicates; representative results are shown.

### Assessing the SY^LL cleavage *in vivo* with a yeast model

To examine ZMPSTE24-dependent cleavage of prelamin A *in vivo*, we expressed human ZMPSTE24 and prelamin A in yeast strains lacking yeast Ste24 and we observed SY^LL cleavage by ZMPSTE24 (*i.e.*, endoproteolytic release of mature lamin A) (Fig 7A), as we previously reported [48]. Notably, we show here that the SY^LL cleavage occurred regardless of whether the prelamin A terminated in CSIM or CVLS, or other CaaX motifs, and regardless of whether Rce1 was present or absent. We used a yeast halo assay to determine if Ste24 is capable of producing mature **a**-factor from **a**-factor constructs terminating in CSIM and CVLS [31, 47]. We observed production of mature bioactive **a**-factor from both CSIM and CVLA constructs in WT and *ste24Δ* yeast (Fig 7B). The production of some **a**-factor in *ste24Δ* yeast, albeit a reduced amount, is likely due to the ability of Axl1 to carry out the final N-terminal processing step in the absence of the Ste24-dependent N-terminal processing step [36]. Notably, in *rce1Δ* yeast, only **a**-factor constructs ending with a *bona fide* CaaX motif (CVIA, CVLS) yielded mature **a**-factor; constructs terminating with CTLM and CSIM (sequence motifs in which the cysteine is followed by a polar residue) did not yield mature **a**-factor (Fig 7B). This observation is likely explained by the fact that **a**-factor secretion and activity require carboxymethylation of a C-terminal farnesylcysteine (which is only possible after a tripeptide cleavage) [36]. The CSIM sequence likely undergoes dipeptide cleavage, thereby preventing carboxymethylation and the production of mature **a**-factor. Our findings suggest that both ZMPSTE24 and Ste24 carry out a tripeptide cleavage when proteins terminate with a *bona fide* CaaX motif but not when proteins terminate with a modified C-terminal sequence motif where the cysteine is followed by a polar residue (*e.g*., Thr, Ser).

**Fig 7.**
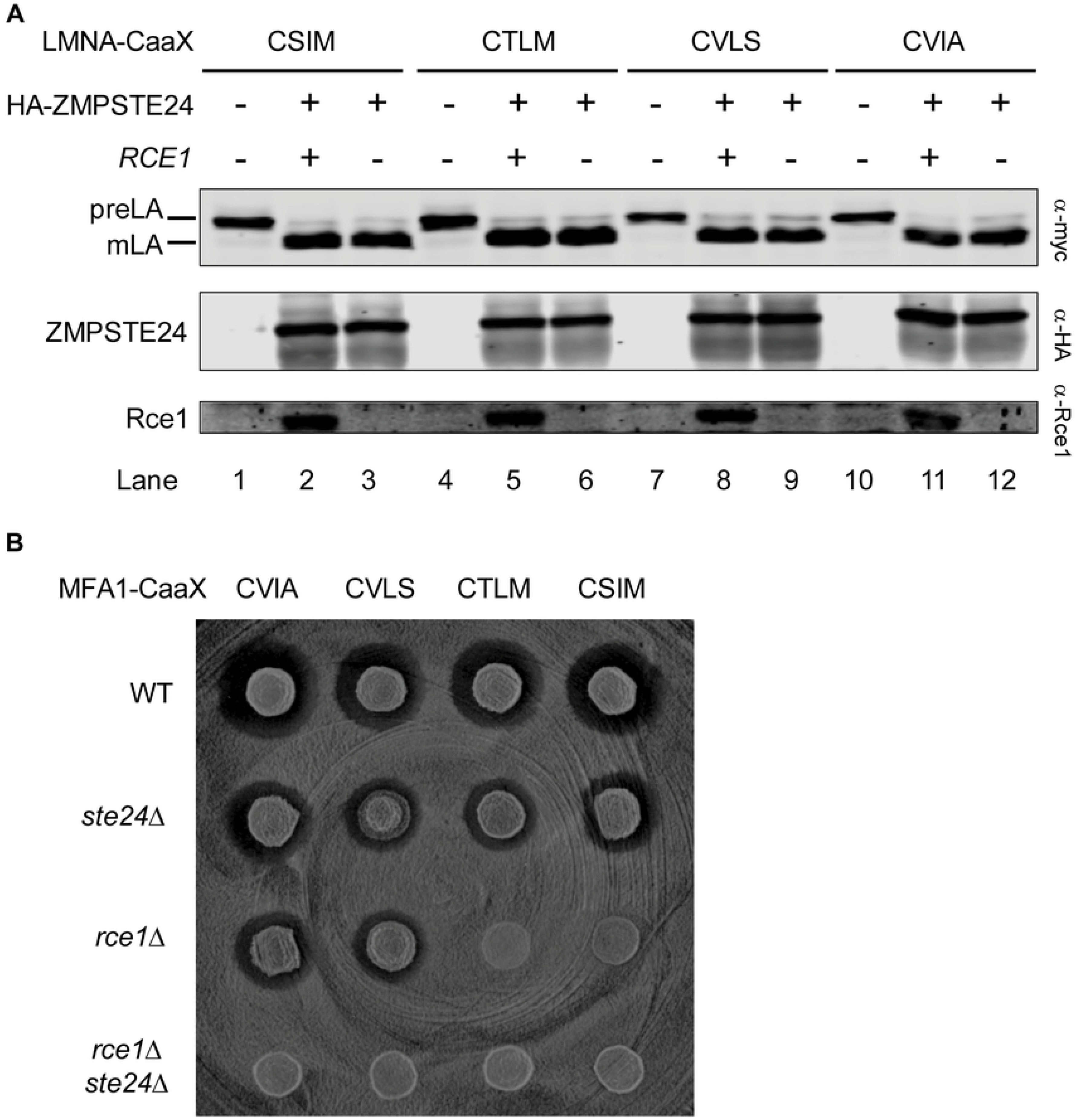
Prelamin A processing by yeast that express both ZMPSTE24 and human prelamin. (A) **(A)** ZMPSTE24-mediated cleavage of myc-tagged prelamin A proteins (wild-type prelamin A, terminating with CSIM, and mutant prelamin A proteins terminating in CVIA, CVLS, or CTLM) by rce1Δste24Δ or ste24Δ yeast. Processing of prelamin A to mature lamin A was assessed by SDS-PAGE and western blotting with a myc antibody. Pre LA, prelamin A; mLA, mature lamin A. (B) “Halo” assay of mature **a**-factor production by different yeast strains that had been transformed with **a**-factor constructs terminating with CVIA, CVLS, CTLM, or CSIM.

## Discussion

Although the requirement for ZMPSTE24 activity in prelamin A processing has been known for some time [1, 2], this is the first study to use human prelamin A peptides to rigorously examine prelamin A processing at a biochemical level. Our results suggest that the often-repeated and well-accepted paradigm for prelamin A processing needs revision (Fig 8A, compared to S1 Fig). First, we show that the rate of cleavage by ZMPSTE24 at the upstream SY^LL site is five times greater than at the CSIM site. Second, we showed that SY^LL cleavage is not dependent on prior CSIM processing. Instead, when both sequences are present in one peptide, ZMPSTE24 first cleaves at the SY^LL site then at the CSIM site (Fig 8A). Hence, the SY^LL cleavage occurs *before* the C-terminal processing. Third, we show that processing of the C-terminal CSIM involves a CS^IM dipeptide cleavage reaction, not a C^SIM tripeptide cleavage. The dipeptide cleavage was observed regardless of peptide length, whether or not the peptide was farnesylated, whether the ZMPSTE24 had been purified from insect or mammalian cells, or whether ZMPSTE24 was in a membrane or a detergent micelle. The discovery of a dipeptide cleavage is consistent with our atomic structure of ZMPSTE24 as a complex with a CSIM tetrapeptide, which revealed that the CSIM peptide was perfectly aligned for a dipeptide cleavage [34] (Fig 9).

**Fig 8.**
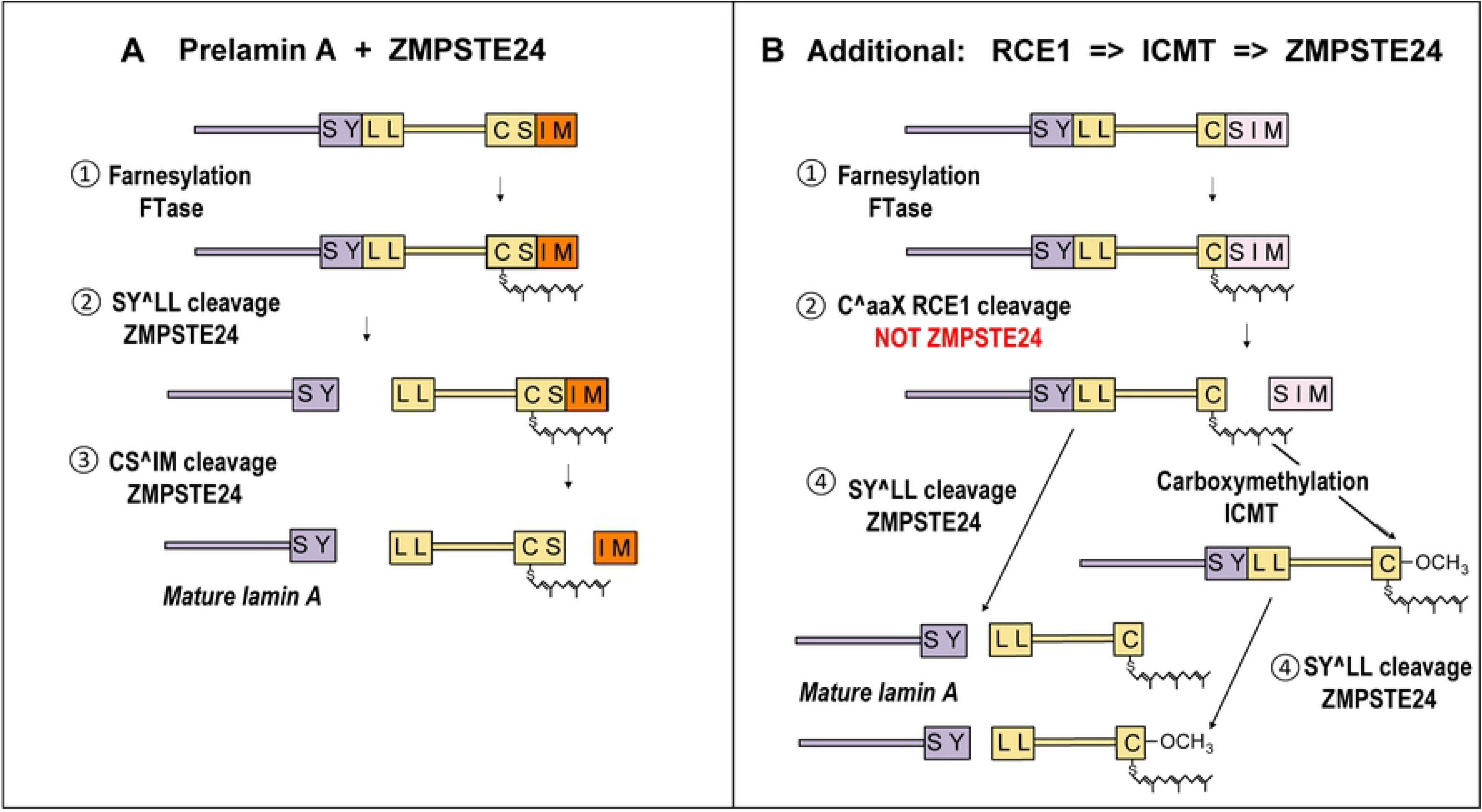
Proposed pathways for ZMPSTE24-mediated prelamin A processing. (A) The “ZMPSTE24-only” processing pathway, in which the SY^LL cleavage occurs before the CS^IM cleavage. (B) Additional potential processing routes when RCE1 and ICMT are present in addition to ZMPSTE24. In this case farnesylated prelamin A could be processed by RCE1-mediated release of the SIM tripeptide from the C-terminal CSIM motif. This product could then undergo SY^LL cleavage by ZMPSTE24 directly or be further modified by ICMT before the SY^LL cleavage.

**Fig 9.**
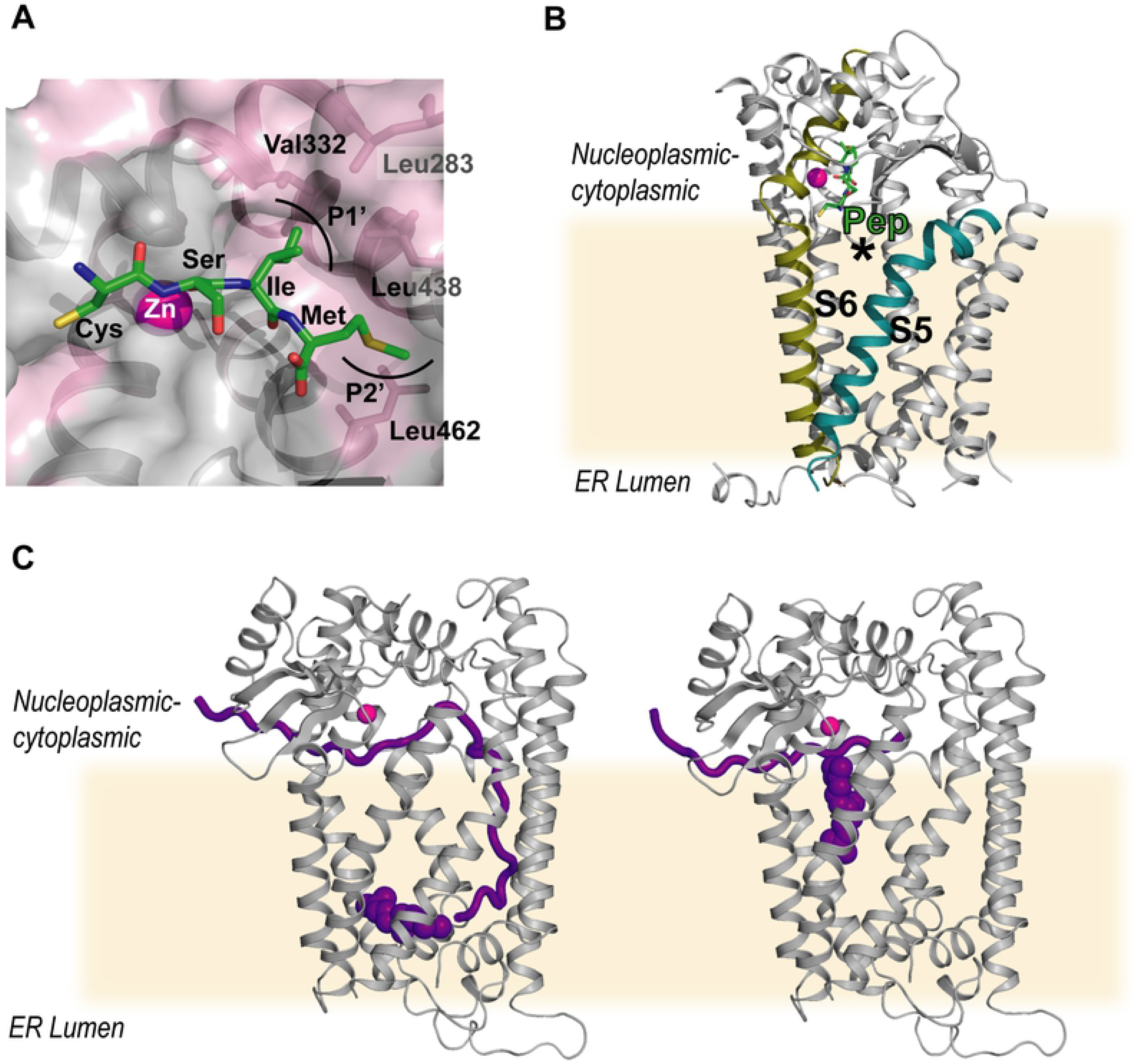
Position of peptides in the ZMPSTE24 structure. (A) Alignment of CSIM peptide in the active site of ZMPSTE24 (PDB: 2YPT). Hydrophobic residues (Ala, Val, Ile, Leu, Phe and Met) are in pink, peptide is shown as stick model, Zn is in magenta, residues involved in the formation of P1’ and P2’ pocket are labelled. (B) Structural overview of ZMPSTE24 and CSIM peptide. Peptide is shown as stick model, Zn is in magenta, S5 and S6 in ZMPSTE24 are highlighted. (C) Model of ZMPSTE24 with two farnesylated peptides resembling the SY^LL (left) and CS^IM (right) cleavage event. Peptides are in purple and the zinc ion is in magenta.

The unexpected dipeptide cleavage may be due to the fact that prelamin A’s C-terminal CSIM sequence is not in fact a classic CaaX motif, in that the serine residue following the cysteine is polar, not aliphatic. When we examined a prelamin A peptide terminating with the canonical CaaX motif found in HRAS (CVLS), ZMPSTE24 carried out the classic C^VLS tripeptide cleavage. The presence of a nonpolar valine after the cysteine in the CVLS sequence facilitates hydrophobic contacts in the P1’ site in ZMPSTE24’s substrate-binding site, which allows alignment for a tripeptide cleavage. Thus, ZMPSTE24 can act as a classic CaaX protease with a typical CaaX motif. However, ZMPSTE24’s only known substrate, prelamin A, does not have a classic CaaX motif. Instead it has a polar serine after the cysteine, and leading to dipeptide cleavage.

We took advantage of yeast expressing both human ZMPSTE24 and full-length human prelamin A constructs to examine ZMPSTE24-mediated prelamin A processing in living cells. Consistent with our biochemical studies with prelamin A peptides, ZMPSTE24 faithfully carried out the upstream SY^LL cleavage within prelamin A, releasing mature lamin A, regardless of whether the prelamin A terminated in CSIM or was modified to terminate with CVIA, CVLS, or CTLM. Yeast **a**-factor biogenesis studies were also consistent with our *in vitro* biochemical findings. The biogenesis, and secretion of mature active **a**-factor, which can be monitored with a halo assay, requires farnesylation, a tripeptide Caa^X cleavage, and methylation of the newly exposed farnesylcysteine, followed by upstream cleavages within the **a**-factor precursor. In our current studies, we showed that Ste24, the yeast orthologue of ZMPSTE24, is capable of promoting **a**-factor biogenesis in *rce1Δ* yeast from **a**-factor precursors that terminate with CVIA or CVLS but not from precursors that terminate with CTLM or CSIM. It is likely ZMPSTE24 carries out a C-terminal dipeptide rather than tripeptide cleavage, in the CTLM and CSIM proteins, preventing farnesylcysteine methylation and thereby preventing secretion of mature active **a**-factor which requires farnesylation and methylation for secretion and activity [36]. In contrast, Ste24 likely carries out a tripeptide cleavage in the CVIA and CVLS **a**-factor precursors, allowing carboxyl methylation of the farnesylcysteine and production of mature **a**-factor. Consistent with this interpretation, mature bioactive **a**-factor was produced from the CTLM and CSIM precursors in yeast that expressed Rce1, which is a *bona fide* CaaX tripeptide cleavage enzyme.

The significance of the farnesylation on the C-terminal CSIM cysteine is an intriguing question. In our biochemical studies, we observed cleavage of nonfarnesylated prelamin A peptide at the SYLL site, suggesting that protein farnesylation is not absolutely critical for processing. However, *in vivo* evidence with genetically modified mice suggests that the SY^LL cleavage occurs only after protein farnesylation. Davies *et al.*, [49] created prelamin A knock-in mice in which the C-terminal CSIM motif was changed to SSIM, thus abolishing prelamin A farnesylation. In these mice, there was no detectable processing of the nonfarnesylated prelamin A to mature lamin A, despite normal ZMPSTE24 expression. In addition, a prelamin A with a C-terminal CSIM sequence synthesized in yeast that are expressing either yeast Ste24 or human ZMPSTE24, the prelamin A is correctly processed. However, prelamin A with a C-terminal SSIM sequence is not cleaved [48]. These findings imply that *in vivo* the SY^LL cleavage depends on prelamin A farnesylation. Farnesylation of prelamin A may facilitate localisation to the membrane surface, bringing it in close proximity to the intramembrane ZMPSTE24.

In cells, the C-terminus of prelamin A will also be exposed to RCE1 and is thus a substrate for RCE1-mediated C^SIM tripeptide cleavage (Fig 8B). However, ZMPSTE24-mediated processing of prelamin A, with release of mature lamin A, does occur in RCE1- or ICMT-deficient fibroblasts, although the efficiency is slightly reduced (evident from a small amount of uncleaved prelamin A in cells) [42]. In any case, it seems likely that there will be a variety of modifications to the C-terminus of prelamin A and that the proportion of each modification will depend on expression and localisation of the substrates, intermediates and enzymes. However, it would appear that ZMPSTE24 is capable of processing farnesylated prelamin A, and any of the C-terminal modified prelamin A variants, removing 15-18 residues, including the farnesyl membrane anchor, thus producing the required mature lamin A.

The physiologic purpose for prelamin A processing, and why prelamin A processing has been conserved through mammalian evolution, has remained elusive. Coffinier *et al.*, [50] generated a knock-in mouse in which a stop codon was introduced into the prelamin A gene after the codon for the last amino acid in mature lamin A. This mouse produced mature lamin A directly (bypassing ZMPSTE24-mediated processing). Surprisingly, no overt histopathology was observed in those mice, but fibroblasts from the mice had greater numbers of nuclear blebs and appeared to have relatively lower amounts of lamin A at the nuclear rim. In light of these findings, we suspect ZMPSTE24-mediated processing of prelamin A optimizes targeting of mature lamin A to the nuclear rim. While direct production of mature lamin A did not result in overt histopathology in laboratory mice, we nevertheless suspect that optimized delivery of mature lamin A is probably important for optimal function of the nuclear lamina and for that reason has been favored during evolution. Conversely, it is clear that that lack of prelamin A processing by ZMPSTE24, with accumulation of permanently farnesylated prelamin A, is detrimental to human health.

Given the importance of prelamin A post-translational processing in a range of conditions such as progeria, lipodystrophies, and potentially even in normal ageing, an understanding of how ZMPSTE24 functions in normal and pathological conditions is essential. This work provides a new paradigm for the role of ZMPSTE24 in prelamin A processing, suggesting alternative views of its activity in cells. Now that we are clear which reaction is central to ZMPSTE24 function, we are in a better position to develop therapies for ZMPSTE24 related diseases.

## Materials and Methods

### Peptides and reagents

The peptide substrates were designed based on sequence of the C terminus of human prelamin A and purchased from commercial sources: Peptides 1 to 4 were purchased from Peptide Protein Research Limited (Fareham, United Kingdom) and Peptide 5 was purchased from Severn Biotech (Kidderminster, United Kingdom). The sequences are shown in Table 1. Nε-DNP-L-lysine hydrochloride, and 2-Aminobenzoic acid were purchased from Sigma-Aldrich Company Ltd. The peptides, Abz, and K(Dnp) were dissolved in DMSO at a final concentration of 10 mM. The stocks were aliquoted and kept at −20°C for future use.

### Protein expression and purification

ZMPSTE24 was expressed and purified as previously published with modifications [34]. In brief, the full-length protein was expressed using Bac-to-Bac® Baculovirus Expression System (Invitrogen) with a C-terminal tobacco etch virus (TEV) cleavage site, a His10 tag and a FLAG tag (pFB-CT10HF-LIC). *Spodoptera frugiperda* (Sf9) cells (Thermo-Fisher Scientific, Cat. No. 11496015) at the density of 2×10^6^ cell per ml were infected with 5 ml of P2 (second passage) recombinant baculovirus in Sf-900 II SFM medium and incubated for 72 h at 27°C. For expression in the HEK293 derived Expi293F™ cells (Thermo-Fisher Scientific, Cat. No. A14527), full-length ZMPSTE24 gene was cloned into the pHTBV1.1-LIC baculovirus transfer vector (The BacMam vector backbone, pHTBV1.1, was kindly provided by Professor Frederick Boyce, Massachusetts General Hospital, Cambridge, MA and adapted for ligation independent cloning in house). This vector provides with a C-terminal TEV cleavage site followed by a His10 tag and a Twin-Strep-tag^®^ (pHTBV1.1-CT10H-SIII-LIC). 1 Litre of Expi293F™ cells at the density of 2×10^6^ cell per ml were infected with 30 ml of P3 (third passage) recombinant baculovirus in Freestyle 293 expression medium and incubated for 48 h at 37 °C, 8 % CO_2_.

All the following steps were performed at 4°C unless otherwise indicated. The cells were resuspended in lysis buffer (50 mM HEPES-NaOH, pH 7.5, 200 mM NaCl, Roche protease inhibitor cocktail) and lysed by two passes through an EmulsiFlex-C5 homogenizer (Aventis). The membrane proteins were extracted from the cell lysate with 1% DDM for 1 h. Cell debris was removed by centrifugation at 35000 g for 1 h. The supernatant was subsequently incubated with Co^2+^ charged TALON resin (Clontech) for 1 h. After washed with 30 column volumes of washing buffer (50 mM HEPES-NaOH, pH 7.5, 200 mM NaCl, 0.03% DDM and 20 mM imidazole-HCl), the target protein was eluted with elution buffer (washing buffer supplemented with 250 mM imidazole-HCl). The eluted protein was concentrated in a 100 kDa cut-off Vivaspin® 2 Centrifugal Concentrator to a final volume of less than 500 μl before injected onto a Superdex 200 10/300 Increase GL column (GE Healthcare), which was equilibrated with GF buffer (20 mM HEPES-NaOH, pH 7.5, 200 mM NaCl, 0.01% DDM) prior to the injection. The peak fractions were pooled together and TEV protease was added at a weight ratio of 1:10 for overnight treatment. The His-tagged TEV protease was removed by a second TALON resin binding for 1 h, and the flow through was collected and analyzed by SDS-PAGE and MSD-ToF electrospray ionization orthogonal time-of-flight mass spectrometer (Agilent Technologies Inc.). The protein was stored at 4°C for up to 2 days or flash-frozen and kept at −80°C for longer periods.

### *In vitro* proteolysis assay

In brief, the 10 mM peptide stock was firstly diluted to 1 mM in 100 mM HEPES-NaOH, pH 7.5 and further diluted to twice of the desired concentrations in GF buffer. The concentration of purified ZMPSTE24 was adjusted to 0.08 mg ml^−1^ (1.4 μM). For peptides 1, and 2, assays were performed in a 384 well microplate (Corning #3573) by mixing 10 μl of the peptide solution and equal volume of the protein solution. For 0 μM peptide concentration, GF buffer containing 2% DMSO was used instead of GF buffer as the control. The fluorescence was monitored using a SpectraMax M2^®^ (Molecular Devices) plate reader at regular intervals over a 30 min time course. The reactions were carried out at 37°C, the excitation and emission wavelengths were set to 320 and 415 nm, respectively. For Peptide 3, instead of a dilution series, the assay was only performed at the peptide concentration of 100 μM as a relative comparison to the cleavage efficiency to Peptide 1. To compare the activity of Expi293™ and Sf9 produced ZMPSTE24, 50 μM Peptide 1 and 100 μM Peptide 2 was used respectively for a time course measurement.

### Activity and kinetic analysis

The raw data were graphed using Microsoft Excel. The relative fluorescent intensity (RFI) was converted to concentration units (nM) by referencing the standard curve, in which the RFI of a 1:1 mixture of fluorophore and quencher pair at varying concentrations (0 to 90 μM) was measured in the presence of the same range of substrate concentrations (0 to 90 μM).

The initial velocities (nM s^−1^) were plotted against the substrate concentrations and analyzed using PRISM software (GraphPad Software Inc.). To obtain the apparent kinetic parameters (*i.e. K_m_* and *V*_max_), the graph was fitted to a non-linear regression one-phase decay equation as the catalytic reaction obeyed Michaelis-Menten kinetics. The S.E. values shown represent error with respect to curve fitting.

### Mass Spectrometry

All the mass spectrometry measurements in this study were performed using an Agilent 1290 Infinity LC System in-line with an Agilent 6530 Accurate-Mass Q-TOF LC/MS (Agilent Technologies Inc.). The reaction mixture of the proteolysis assay at 100 μM peptide concentration was used as the sample for mass spectrometry analysis. 5 μl of the reaction mixture was diluted to 60 μl with 30% methanol in 0.1% formic acid. 60 μl of sample was injected onto a ZORBAX StableBond 300 C3 column (Agilent Technologies Inc.) by an auto sampler. The solvent system consisted of 0.1% formic acid in ultra-high purity water (Millipore) (solvent A) and 0.1% formic acid in 100% methanol (solvent B). Initially, 30% solvent B was applied at a flowrate of 0.5 ml/min. The sample was eluted by a linear gradient from 30% to 95% of solvent B over 7 min. Elution was then isocratic at 95 % B for 2 min, followed by a further 2 min equilibration at 30% B. The mass spectrometer was operated in positive ion, 2 GHz detector mode. Source parameters were drying gas 350 °C, flow 12 l/min, nebulizer 60 psi, capillary 4000 V. Fragmentor was 250 V, collision energy 0 V and data acquired from 100-3200 m/z. Protein or peptide (fragments of peptide) intact mass was acquired using an MSD-ToF electrospray ionisation orthogonal time-of-flight mass spectrometer (Agilent Technologies Inc.). Data analysis was performed using MassHunter Qualitative Analysis Version B.07.00 (Agilent) software.

### Time course mass spectrometry

The time course mass spectrometry was performed to characterize the cleavage of Peptide 4, 5 and 6. In brief, the peptide stock was prepared as previously specified. 20 μl of WT ZMPSTE24 at the concentration of 0.08 mg ml^−1^ was mixed with 20 μl of 200 μM Peptide 4, 5 or 6, and incubated at 37°C in a heating block. 5 μl of the reaction mixture was taken out and added into 55 μl of 30% methanol to terminate the reaction at time point 1 min, 2 min, 5 min, 10 min, 30 min and 60 min. The samples were kept on ice until loaded onto the Q-TOF LC/MS. The mass spectrometry analysis was performed as previously described. When looking for specific species from the reaction mixture, the extracted ion chromatogram (EIC) function was used.

### *In vivo* yeast prelamin A cleavage assay

*In vivo* cleavage reactions of myc-tagged LMNA_431-661_-CaaX (herein designated myc-LMNA-CaaX) were performed using strains SM5877 (*rce1Δste24Δ*), SM6303 (*ste24Δ HA-ZMPSTE24*) and SM6437 (*rce1Δste24Δ HA-ZMPSTE24*) transformed with plasmids pSM3371 (myc-LMNA-CSIM), pSM3372 (myc-LMNA-CTLM), pSM3458 (myc-LMNA-CVLS) or pSM3511 (myc-LMNA-CVIA)[48]. These yeast strains were grown in SC-leu medium to OD_600_ 1.5-2, collected by centrifugation, and processed for SDS-PAGE [51]. Lysates (0.3 OD_600_ cell equivalents) were resolved by 10% SDS-PAGE, proteins were transferred to nitrocellulose (Bio-Rad Trans-Blot^®^ Turbo™) and the membrane blocked using Western Blocking Reagent (Roche). Lamin proteins were detected using mouse anti-myc antibodies (clone 4A6, Millipore cat #05-724; 1:10000 dilution) decorated with goat anti-mouse secondary IRDye 680RD antibodies (LI-COR). Blots were re-probed to detect ZMPSTE24 and Rce1 using rat anti-HA (clone 3F10, Roche cat #11867423001; 1:10000 dilution), and polyclonal rabbit anti-Rce1 [52] (1:2000 dilution). LI-COR secondary antibodies used were goat anti-rat IRDye 680RD and goat anti-rabbit IRDye 800CW, respectively.

The halo assay was performed using yeast strains SM2331 (*mfa1Δmfa2Δ*), SM3375 (*mfa1Δmfa2Δste24Δ*), SM3689 (*mfa1Δmfa2Δrce1Δ*) and SM3691 (*mfa1Δmfa2Δste24Δrce1Δ*) transformed with *MFA1-CaaX* plasmids pSM478 (CVIA), pSM3500 (CVLS), pSM3509 (CTLM) and pSM3498 (CSIM) [36]. These strains were grown in SC-Ura medium, concentrated by centrifugation and spotted onto a lawn of SM2375 (MAT α *sst2*) cells plated on YPD + 0.4% Triton X-100. Plates were incubated for 2 days at 30°C and scanned.

### Proteoliposome reconstitution

A stock of phosphatidyl-choline (POPC) and phosphatidyl-ethanolamine (POPE) (Avanti) at 3:1 (w/w) was dried under argon and re suspended in detergent free GF buffer to a final concentration of 20 mg ml^−1^. The liposomes were disrupted by the addition of 22 mM sodium cholate (Anatrace). Purified ZMPSTE24 was added into the mixture at lipid protein ratio (LPR) at 20:1 (w/w). After 30 min incubation on ice, Bio-Beads SM-2 (Bio-rad) were supplemented into the mixture to remove the detergent. The Biobeads were changed to fresh batch after 2 h for an overnight incubation. A further 1 h incubation with fresh Biobeads was performed the next morning. To make sure the complete removal of detergent, the proteoliposome was centrifuged at 100, 000 g for 1 h and the pellet was re-suspended in detergent free GF buffer at lipid concentration 5 mg/ml. The resuspension was then applied onto a LiposoFast extruder (avestin) and extruded against a membrane with a pore size of 400 nm.

## Abbreviations

DDM: n-dodecyl β-D-maltoside
FRET: fluorescence resonance energy transfer
GFP: green fluorescent protein
HGPS: Hutchinson-Gilford progeria syndrome
MAD-B: mandibuloacral dysplasia type B
RD: restrictive dermopathy
Abz: 2-Aminobenzoyl
Dnp: 2, 4-dinitrophenyl
RFI: relative fluorescence intensity

## Acknowledgements

We thank Frederick Boyce (Massachusetts General Hospital) for providing the BacMam vector (pHTBV1.1) with VSV-G and enhanced CMV promoter, which was adapted for ligation independent cloning by the SGC BioTech group. The authors declare no competing financial interests.

## Funding

This work was funded by an Medical Research Council (MRC) grant no MR/L017458/1 to EPC, an National Institute of Health (NIH) grant to SM (5R35GM127073) and a National Heart, Lung and Blood Institute (NHLBI) grant to SY (HL139725). LN, AQ, YYD, AC, RC, SMMM, NBB, ACWP and EPC is funded by the Structural Genomics Consortium (SGC). The SGC is a registered charity (number 1097737) that receives funds from AbbVie, Bayer Pharma AG, Boehringer Ingelheim, the Canada Foundation for Innovation, Genome Canada, Janssen, Lilly Canada, Merck KGaA, Merck & Co., Novartis, the Ontario Ministry of Economic Development and Innovation, Pfizer, São Paulo Research Foundation-FAPESP and Takeda, as well as the Innovative Medicines Initiative Joint Undertaking ULTRA-DD grant 115766 and the Wellcome Trust (106169/Z/14/Z). The authors declare no competing financial interests.

